# CXCR4 and MIF are required for neutrophil extracellular trap release triggered by *Plasmodium*-infected erythrocytes

**DOI:** 10.1101/852574

**Authors:** Danielle S. A. Rodrigues, Elisa B. Prestes, Leandro de Souza Silva, Ana Acácia S. Pinheiro, Jose Marcos C. Ribeiro, Alassane Dicko, Patrick E. Duffy, Michal Fried, Ivo M. B. Francischetti, Elvira M. Saraiva, Heitor A. Paula Neto, Marcelo T. Bozza

## Abstract

Neutrophil extracellular traps (NETs) evolved as a unique effector mechanism contributing to resistance against infection that can also promote tissue damage in inflammatory conditions. Malaria infection can trigger NET release, but the mechanisms and consequences of NET formation in this context remain poorly characterized. Here we show, similarly to previous reports, that patients suffering from severe malaria had increased amounts of circulating DNA and increased neutrophil elastase (NE) levels in plasma. We used cultured erythrocytes and isolated human neutrophils to show that *Plasmodium*-infected red blood cells release MIF, which in turn caused NET formation by neutrophils in a mechanism dependent on the C-X-C chemokine receptor type 4 (CXCR4). NET production was dependent on histone citrulination by PAD4 and independent of reactive oxygen species (ROS), myeloperoxidase (MPO) or NE. In vitro, NETs functioned to restrain parasite dissemination in a mechanism dependent on MPO and NE activities. Finally, C57/B6 mice infected with *P. berghei* ANKA, a well-established model of cerebral malaria, presented high amounts of circulating DNA, while treatment with DNAse increased parasitemia and accelerated mortality, indicating a role for NETs in resistance against *Plasmodium* infection.

**Author summary:** Protozoans of the Plasmodium genre infect red blood cells and cause malaria in humans and various other mammalian species. Estimated malaria cases are at more than 200 million, with 450,000 deaths per year, being cerebral malaria a serious complication that accounts for the majority of deaths. Neutrophils are cells that participate in host defense against pathogens. These cells use various mechanisms to kill invading microrganisms, including the release of webs of DNA, called neutrophil extracellular traps (NETs). These NETs can help control infections but can also induce tissue damage and their role in malaria and the mechanisms of NET production during malaria infection are starting to be understood. Here we show that infected red blood cells produce a cytokine, macrophage migration inhibitory factor (MIF) that stimulates neutrophils to release NETs. These NETs function to limit *Plasmodium* dissemination and, thus, digestion of NETs with DNAse treatment causes increased parasitemia and accelerated death in an experimental model of cerebral malaria. Our study uncovers the mechanism by which infected red blood cells stimulate neutrophils to release NETs and suggest an important participation of this process in malaria control.

## Introduction

Malaria is a highly prevalent and widespread infectious disease caused by protozoans of the *Plasmodium* genus. Amongst the known agents of human malaria, *Plasmodium falciparum* is associated with the complicated forms of disease, including the potentially fatal cerebral malaria [1, 2]. Severe forms of malaria infection can be associated with either impaired mechanisms of resistance - and consequently high parasitemia [3, 4] - or exacerbated tissue damage due to ineffective mechanisms of disease tolerance [5–8]. Studies on the immunological mechanisms of tissue injury and host resistance to malarial infection have generally focused on adaptive immune responses coordinated by CD4+ T cells through the activation of CD8+ T and B cells [9–11]. However, mounting evidences, from both human studies and the *P. berghei* ANKA murine model of severe malaria, point to the involvement of other cell types, including platelets, macrophages and neutrophils [12–15].

Neutrophils participate in the immune response to pathogens by using several mechanisms of killing, including reactive oxygen species (ROS) production, phagocytosis, and the release of antimicrobial peptides and cytotoxic enzymes [16]. Neutrophils are capable of phagocytosing opsonized *P. falciparum* merozoites [17] and *P. falciparum*-infected red blood cells [18]. Neutrophil ROS production positively correlated with *P. falciparum* clearance and individuals with higher ROS production presented faster parasite clearance time (PCT) [19]. These observations would support a beneficial role of neutrophils in mediating *Plasmodium* clearance and disease resistance in malaria. However, in human malaria there is a strong correlation between neutrophil activation markers and disease severity [14, 20, 21], suggesting that overt neutrophil activation may contribute to disease pathogenesis. In fact, neutrophil depletion resulted in decreased brain microhaemorrhages and monocyte sequestration, preventing cerebral malaria development in mice [22, 23]. Therefore, whether neutrophils are beneficial, contributing to pathogen clearance or detrimental, inducing tissue damage during severe malaria remains unresolved.

Neutrophils can release DNA to the extracellular space, which has been shown to function as traps for many different pathogens, including bacteria, fungi, viruses and protozoans. These neutrophil extracellular traps (NET) evolved as a unique innate immune defense mechanism capable of restraining pathogens, avoiding their dissemination and contributing to pathogen killing [24]. However, NETs have also the potential to harm surrounding heathy tissue, thus contributing to both aspects of disease tolerance, i.e. pathogen elimination and collateral tissue damage. It is therefore reasonable to hypothesize that neutrophils and NETs may be involved in malaria pathogenesis. In fact, reports show evidences of NET production in samples from human malaria patients [25, 26]. Moreover, it was recently shown that *Plasmodium*-infected red blood cells are capable of triggering NET release [27]. NET disruption with DNAse treatment resulted in milder lung injury and increased survival in a model of *Plasmodium*-induced acute lung injury [27]. However, the mechanisms involved in NET release in response to *Plasmodium*-infected erythrocytes remain uncharacterized.

In the present study, we show that NETs are released by neutrophils exposed to *Plasmodium*-infected erythrocytes and contribute to restrain pathogen spread and control malaria infection. We also provide evidences that stimulation of NET release is independent of cell-cell contact and is mediated by macrophage migration inhibitory factor (MIF) activation of CXCR4.

## Results

### *P. falciparum*-infected erythrocytes induce NETs

Evidences in both humans and mice suggest that malaria infection triggers NET release. We analyzed blood samples from patients with severe malaria (S.M.) or uncomplicated malaria (U.M.) for signs of NET. We observed that patients with S.M. showed increased circulating neutrophil elastase (NE) levels (Fig 1A) as well as increased circulating nucleosomes (Fig 1B), suggestive of NETs. To further evaluate the potential of *P. falciparum*-infected red blood cells (iRBCs) in inducing NET release, human peripheral blood neutrophils were incubated with infected erythrocytes at increasing neutrophil to iRBC ratios. After 3 h, a significant increase in extracellular DNA content could be detected in supernatant of neutrophils cultured in the presence of infected erythrocytes relative to unstimulated neutrophil controls (Fig 1C). Data representation as absolute extracellular DNA levels showed similar trends (S1A Fig). Increased extracellular DNA content was evident in a 1:1 ratio and was even more pronounced at a 5:1 ratio, reaching a 6-to 7-fold increase relative to control neutrophils (Fig 1C). Incubation of human neutrophils at a lower (0.5:1) erythrocyte:neutrophil ratio did not induce any significant increase in extracellular DNA signal, as well as the incubation with uninfected red blood cells of the same donor at any of the tested ratios.

**Fig 1.**
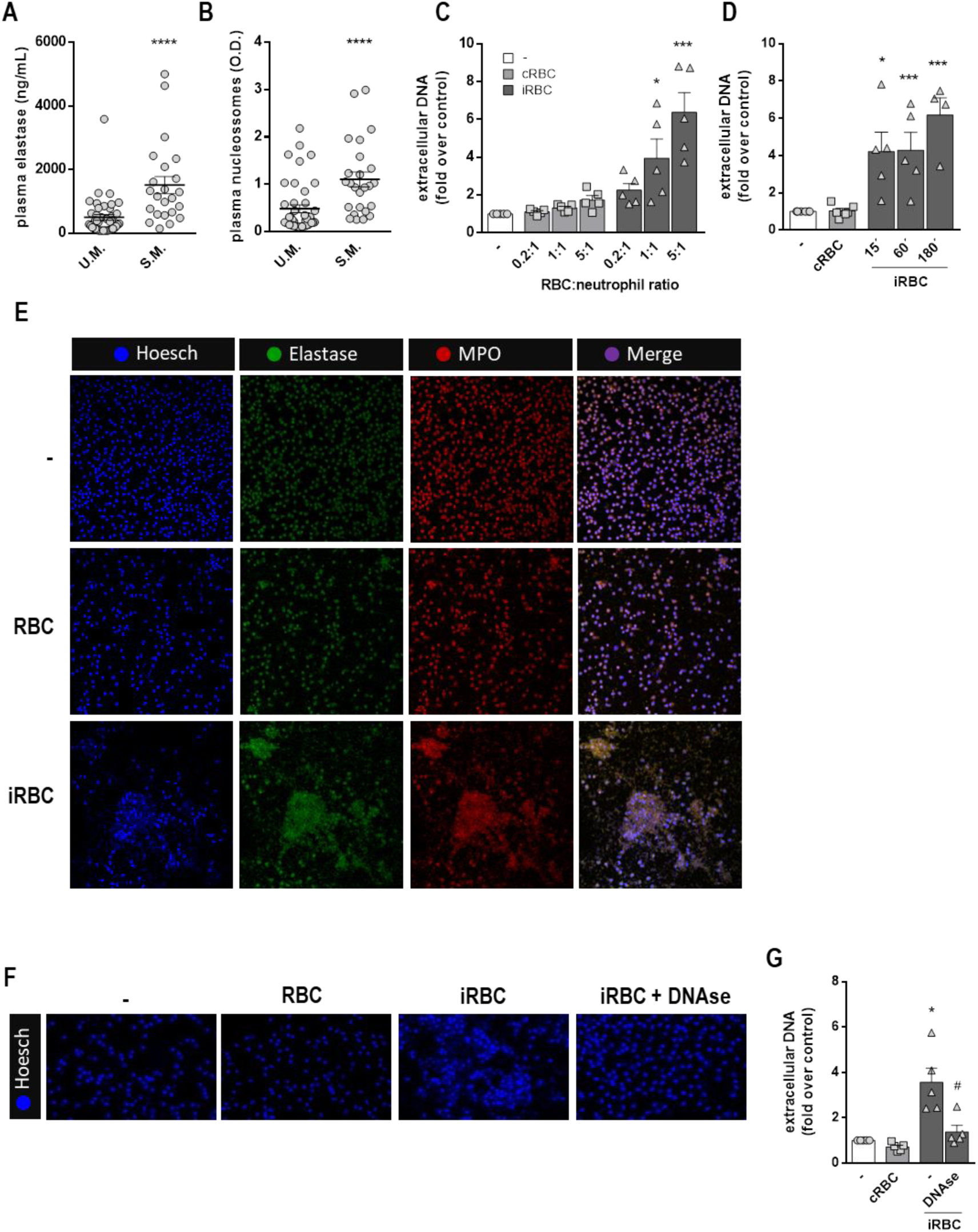
*P. falciparum-*infected erythrocytes induce NETs. (A) Neutrophil elastase and circulating nucleosomes (B) in plasma from human patients diagnosed with uncomplicated malaria (U.M., n=42) or severe malaria (S.M., n=23). Mann-Whitney was performed for statistical significance between groups. P values are indicated in each graph. (C) Fluorimetric determination of NET production by human neutrophils in the presence of *P. falciparum*-infected red blood cells (iRBC) or uninfected RBC (cRBC) at varying red blood cell:neutrophil ratios. (D) Fluorimetric determination of NET production by human neutrophils in the presence of *P. falciparum*-infected red blood cells (iRBC) or uninfected RBC (cRBC) at different time-points. Data in C and D are presented as means ± S.E.M. of the fold induction of extracellular DNA signal relative to resting neutrophils. (E) Visualization by fluorescence microscopy of NETs produced by human neutrophils in the presence of *P. falciparum*-infected (iRBC) or uninfected (cRBC) red blood cells for 3 hours. DNA is stained in blue (Hoesch), neutrophil elastase is stained in green (elastase) and myeloperoxidase is stained in red (MPO). Unstimulated human neutrophils were used as controls. (F) Representative images of the effect of DNAse treatment on NET signal as visualized by fluorescence microscopy. Human neutrophils were incubated with iRBC or cRBC for 3 hours in the presence of DNAse. DNA was stained with Hoesch. (G) Quantification of data derived from (F). Data are presented as means ± S.E.M. of the fold induction of extracellular DNA signal relative to resting neutrophils. * P< 0.05 and *** P<0.001 relative to controls incubated with cRBC, # P< 0.01 relative to untreated control.

NET production induced by infected erythrocytes could be observed at very early time-points, with significant increases being detected at 15 min (Fig 1D). NET production was still high at 60 min and increased further at 180 min (Fig 1D). NET release was paralleled by an increase in lactate dehydrogenase (LDH) activity in culture supernatants (S1B Fig). This suggests that NET production in response to infected erythrocytes is accompanied by cell death, although it is difficult to ascertain whether LDH is derived from NETosing neutrophils or rupturing erythrocytes. We further observed that NET induced by *P. falciparum*-infected erythrocytes showed a cloud-like morphology (Fig 1E). Although we detected a significant extracellular DNA signal using fluorimetric assay as early as 15 minutes, NET-like structures only started to be detectable by immunofluorescence at 60 minutes (data not shown) and peaked at 180 minutes (Fig 1E). NETs stained positively for NE and myeloperoxidase (MPO), two characteristic enzymes found associated to DNA in NETs (Fig 1E). Finally, these NET-like structures, as well as the fluorimetric signal were lost after DNAse incubation (Fig 1F and 1G), suggesting that the structures we are describing here meet the criteria to be classified as NETs. Altogether, these results demonstrate that *P. falciparum*-infected erythrocytes stimulate human neutrophils to release NETs *in vitro*.

### Mechanisms underlying NET production induced by infected erythrocytes

*P. falciparum*-infected erythrocytes triggered a strong ROS production by human neutrophils (S2A Fig). However NET release in response to infected erythrocytes was not inhibited by neither DPI treatment (Fig 2A) nor NAC (S2B Fig), despite their capacity to block ROS production induced by infected erythrocytes (S2C and S2D Figs). These results suggest that NET release in response to infected red blood cells is ROS-independent. Moreover, we observed that uninfected erythrocytes were also able to induce ROS production by human neutrophils, although to a smaller extent (S2C and S2D Figs). This also argues against a possible involvement of ROS in NET release induced by infected erythrocytes since we did not observe any NET production in response to uninfected red blood cells (Fig 1C). MPO and NE were reported to be essential to NET production induced by different stimuli [28, 29]. We used two well described inhibitors of MPO and NE to evaluate the involvement of these two enzymes in NET production induced by *P. falciparum*-infected erythrocytes. Neither inhibitor had any effects on NET production in this model, ruling out the involvement of MPO and NE in this process (Figs 2B and 2C).

**Fig 2.**
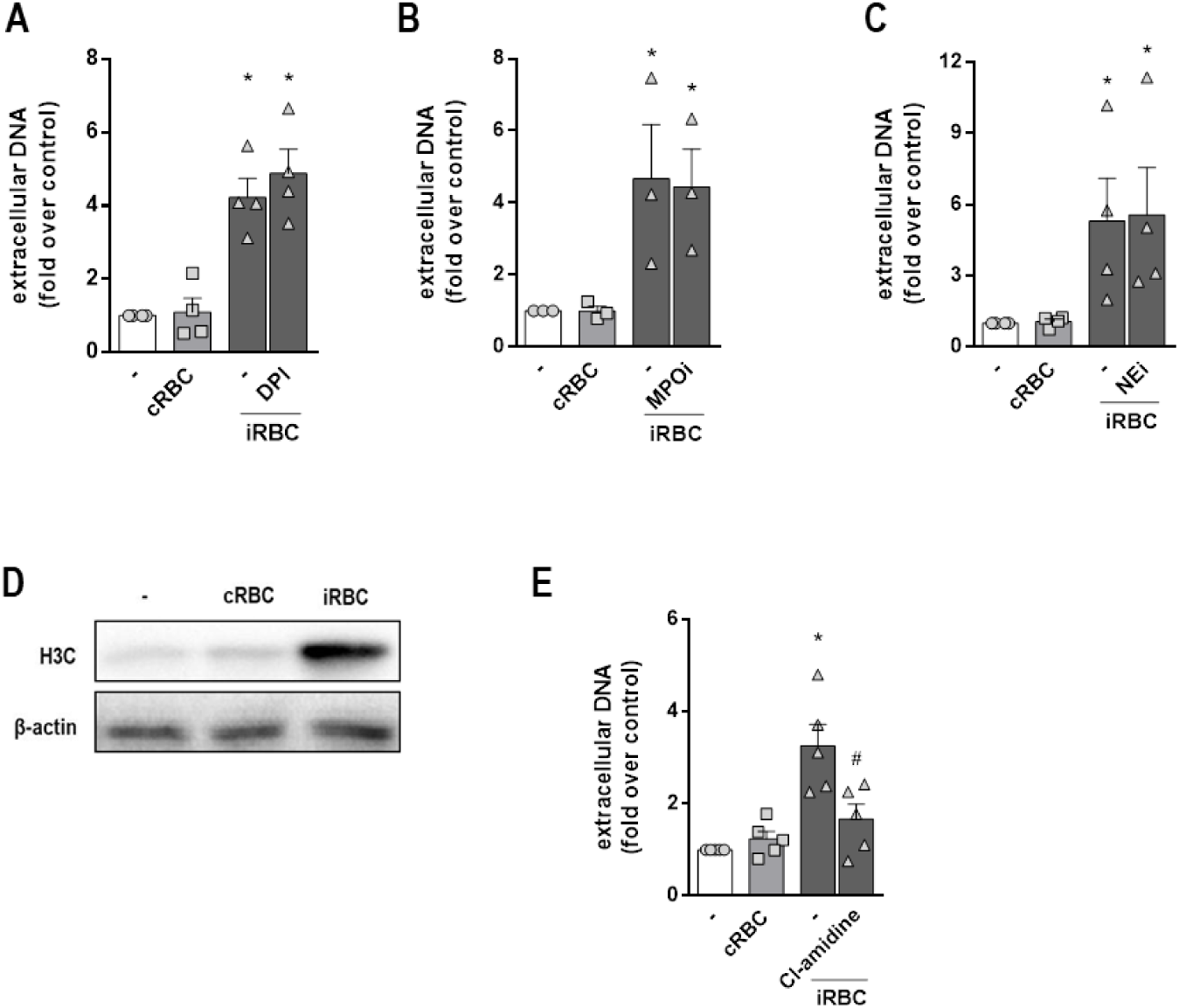
Involvement of ROS, MPO, NE and PAD4 on NET production in response to infected erythrocytes. Human neutrophils were treated with DPI (10 µg/mL) (A), MPO inhibitor (MPOi, 10 µg/mL) (B), neutrophil elastase inhibitor (NEi, 10 µg/mL) (C) or Cl-amidine (12 µM) (E) for 30 minutes and then incubated with *P. falciparum*-infected red blood cells (iRBC) for 3 hours. NET production was determined by fluorimetry. Uninfected red blood cells (cRBC) were used as control. Data are presented as means ± S.E.M. of the fold induction of extracellular DNA signal relative to resting neutrophils. (D) Representative westernblot image of citrullinated histone H3 in extracts of human neutrophils incubated for 3 hours in the presence of infected (iRBC) or uninfected (cRBC) red blood cells. β-actin was used as loading control. * P< 0.05 relative to controls incubated with cRBC, # P< 0.01 relative to untreated control.

Another important step in NET release is histone citrullination by PAD4 [30, 31]. Incubation of human neutrophils with infected erythrocytes induced a strong increase in histone citrullination, as observed by both Western blot (Fig 2D) and immunofluorescence (S3 Fig). Treatment of neutrophils with the PAD4 inhibitor, Cl-amidine, resulted in significant inhibition of NET production (Fig 2E), suggesting the involvement of PAD4-induced histone citrullination in this process. We treated neutrophils with different kinase inhibitors to define the signaling pathways contributing to NET release in response to infected erythrocytes. Incubation of human neutrophils with *P. falciparum*-infected red blood cells induced increased PKCδ expression, in agreement with the observed increased ROS production. We also observed increased phosphorylation of Akt, JNK and p38 (S4A Fig). Inhibition of JNK phosphorylation with SP600125 significantly inhibited NET release (S4B Fig). On the other hand, inhibition of p38 did not have any effect (S4C Fig). Together, our results suggest that NET release by human neutrophils in response to *P. falciparum*-infected red blood cells is dependent on JNK and PAD4, but independent of ROS, NE, MPO and p38.

### NET production induced by infected erythrocytes does not depend on integrins or CD36

Integrins are expressed by neutrophils and mediate a series of their biological functions. Previous studies have implicated CD18/CD11b (Mac-1) in NET production by different stimuli, including *Candida albicans* β-glucan [32] and immobilized immune complexes [33]. We incubated neutrophils with a CD18 blocking antibody during the interaction with infected erythrocytes. We found no effects of anti-CD18, or the isotype control antibody, on NET release induced by infected erythrocytes (Fig 3A). *P. falciparum*-infected erythrocytes interact with endothelial cells through *P. falciparum* erythrocyte membrane protein 1 (PfEMP-1) expressed by infected erythrocytes. PfEMP-1 mediates cytoadhesion of infected erythrocytes to endothelia through its interaction with CD36 and ICAM-1 expressed by endothelial cells [34, 35]. We reasoned that CD36 or ICAM-1 could have a role in mediating the recognition of infected erythrocytes and NET production by human neutrophils. Incubation of neutrophils with an ICAM-1 blocking antibody did not interfere with NET production induced by infected erythrocytes (Fig 3B). Similarly, incubation with an anti-CD36 blocking antibody did not inhibit NET release (Fig 3C). These results show that neither of the integrins known to be involved in NET production or erythrocyte cytoadherence, nor CD36 are involved in the stimulation of NET release by infected erythrocytes. We further treated neutrophils with cytochalasin D, a disruptor of actin polymerization that inhibits phagocytosis. There are evidences of neutrophil phagocytosis of infected red blood cells in human patients and also there are evidences that phagocytosis may inhibit NET release [36]. However, despite these evidences, NET release was neither increased nor decreased by cytochalasin D (Fig 3D), ruling out the involvement of phagocytosis in this process.

**Fig 3.**
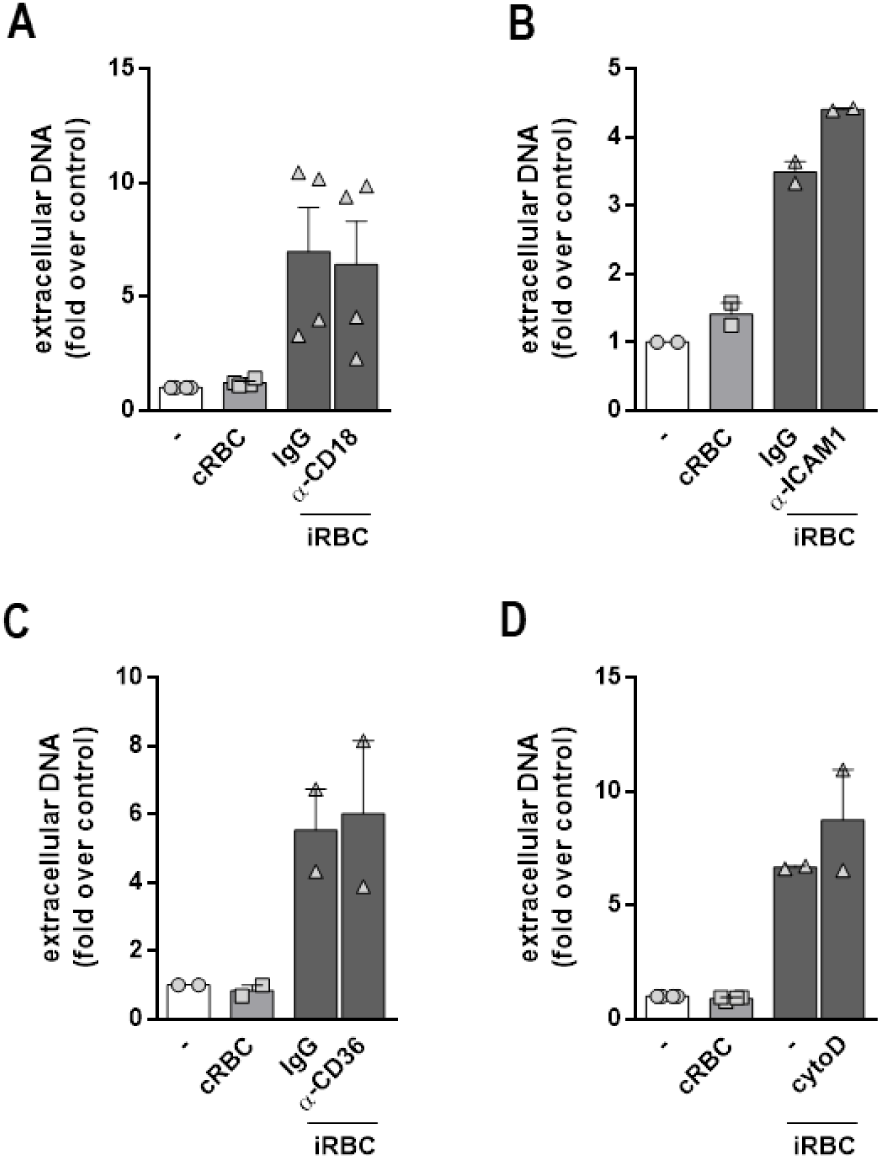
Involvement of CD18, ICAM-1, CD36 and phagocytosis on NET production in response to infected erythrocytes. Human neutrophils were treated with neutralizing antibodies to CD18 (20μg/mL) (A), ICAM-1 (20μg/mL) (B) or CD36 (20μg/mL) (C) for 30 minutes and then incubated with *P. falciparum*-infected red blood cells (iRBC) at a 1:5 ratio for 3 hours. (D) Human neutrophils were treated with cytochalasin D (8 μM) for 30 minutes and then incubated with *P. falciparum*-infected red blood cells (iRBC) at a 1:5 ratio for 3 hours. NET production was determined by fluorimetry. Uninfected red blood cells (cRBC) were used as control. Data are presented as means ± S.E.M. of the fold induction of extracellular DNA signal relative to resting neutrophils. * P< 0.05 relative to controls incubated with cRBC.

### NET production in response to infected RBCs is triggered by macrophage migration inhibitory factor (MIF)

Recently it was demonstrated that aged neutrophils presenting increased CXCR4 expression show enhanced capacity of NET release [37]. Moreover, a report by Sercundes and cols. showed that NETs are involved in pulmonary injury in a murine model of malaria and that CXCR4 inhibition protected mice from acute lung injury [27]. We therefore used AMD3100, a CXCR4 antagonist, to evaluate the involvement of CXCR4 in this model. We observed that AMD3100 inhibited NET release induced by infected red blood cells (Fig 4A). CXCL12 is the typical CXCR4 ligand, but this receptor can also be activated by MIF, which functions as a non-cognate ligand [38]. Moreover, it has been demonstrated that red blood cells are an important source of MIF, contributing with ∼99% of total MIF content in blood [39]. In fact, the cell-permeable MIF antagonist, ISO 1, was able to inhibit NET release in response to infected erythrocytes (Fig 4B). Additionally, treatment with an anti-MIF blocking antibody resulted in a significant inhibition of NET release induced by infected erythrocytes (Fig 4C). Finally, we could detect the presence of MIF in the supernatant of infected, but not uninfected, erythrocytes (Fig 4D). Supernatant derived from cultures of infected erythrocytes induced NET release by human neutrophils (Fig 4E), an effect that could be blocked by anti-MIF antibody (Fig 4F). Altogether, these results indicate that MIF is a soluble mediator released by *P. falciparum*-infected erythrocytes that induce NET release by human neutrophils.

**Fig 4.**
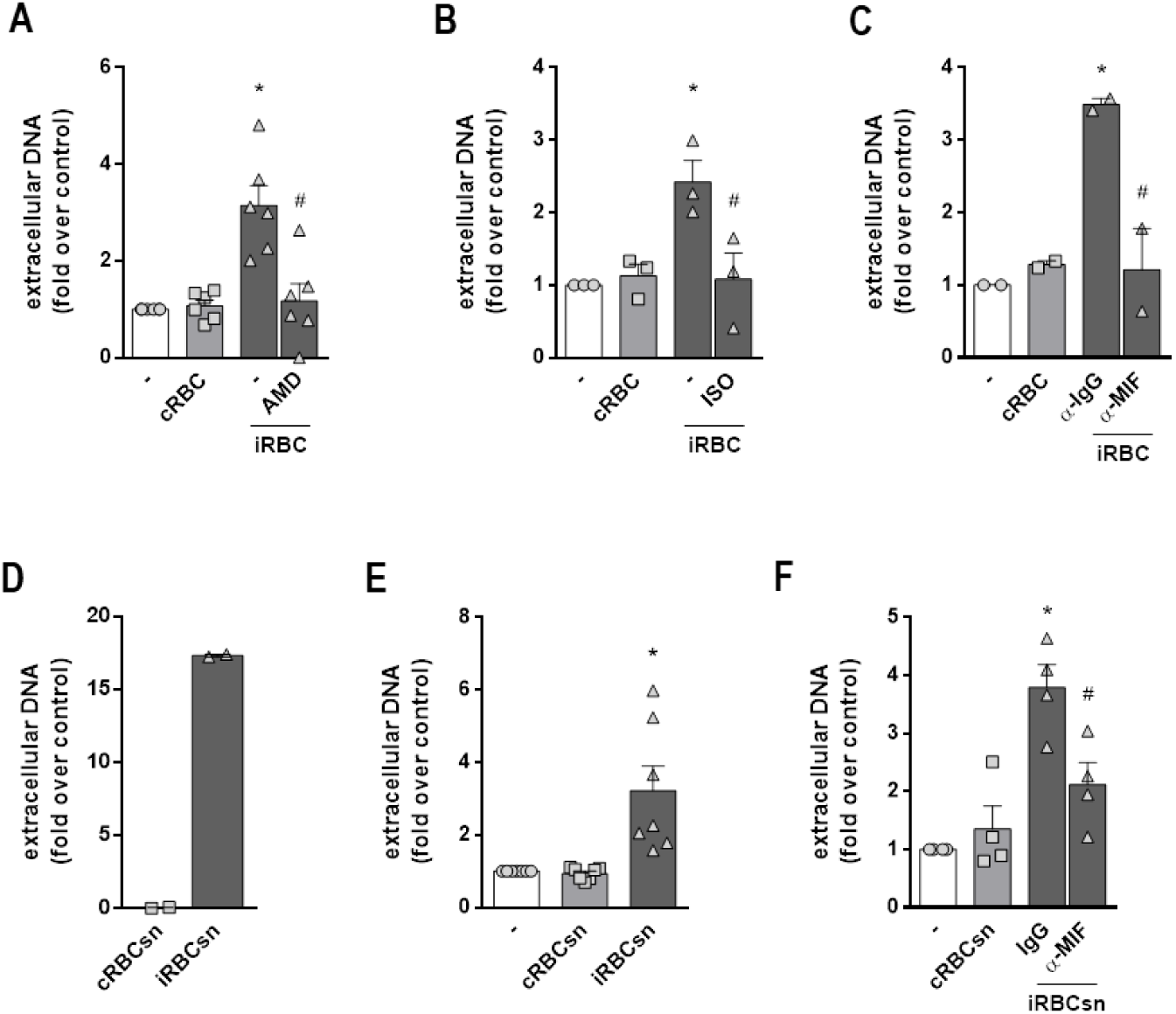
Involvement of CXCR4-MIF axis on NET production in response to infected RBC. Human neutrophils were treated with AMD3100 (AMD, 100 ng/ml) (A), ISO-1 (ISO, 50μM) (B) or a neutralizing anti-MIF antibody (α-MIF, 20μg/mL) (C) for 30 minutes and then incubated with *P. falciparum*-infected red blood cells (iRBC) at a 1:5 ratio for 3 hours. NET production was determined by fluorimetry. Uninfected red blood cells (cRBC) were used as control. Data are presented as means ± S.E.M. of the fold induction of extracellular DNA signal relative to resting neutrophils. (D) Quantification of MIF levels on supernatants from *P. falciparum*-infected (iRBCsn) or uninfected (cRBCsn) red blood cells. (E) Human neutrophils were incubated with supernatants from *P. falciparum*-infected (iRBCsn) or uninfected (cRBCsn) red blood cells and NET production was determined by fluorimetry. (F) Human neutrophils were treated with neutralizing anti-MIF (α-MIF, 20μg/mL) or the appropriate isotype control (IgG) antibody and then incubated with supernatant from *P. falciparum*-infected (iRBCsn) or uninfected red blood cells (cRBCsn). NET production was determined by fluorimetry. Data are presented as means ± S.E.M. of the fold induction of extracellular DNA signal relative to resting neutrophils. * P< 0.05 relative to controls incubated with cRBC, # P< 0.01 relative to untreated control.

### NET restricts parasite dissemination and contributes to host survival

To test the biological significance of NET formation to malaria pathogenesis, we first treated *P. falciparum*-infected erythrocyte cultures with NET rich supernatant collected from human neutrophils previously stimulated with infected erythrocytes. Presence of NETs resulted in fewer ring structures and decreased proportions of infected erythrocytes in culture (Figs 5A and 5B, respectively). Accordingly, DNAse treatment restored the percentage of ring structures to those found in untreated cultures (Fig 5C), suggesting that NETs interfere with *P. falciparum* dissemination *in vitro*. Additionally, treatment of cultures with either MPO or NE inhibitors also resulted in increased levels of ring structures compared to cultures in the presence of NET (Figs 5D and 5E, respectively). This suggests that, despite being dispensable to NET release, MPO and NE activity participate in NET-mediated control of parasite dissemination.

**Fig 5.**
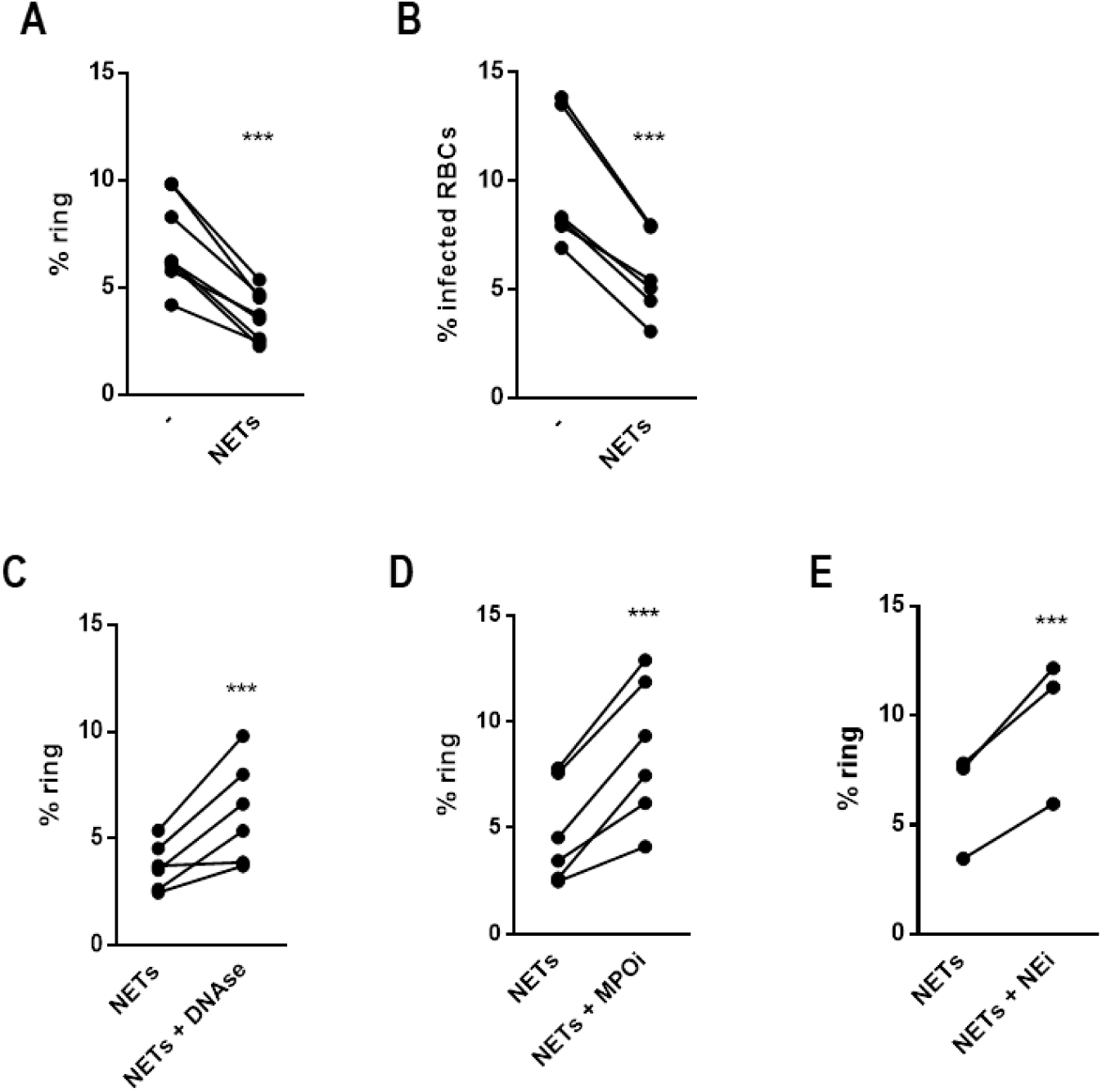
Effect of NETs on *P. falciparum* dissemination in cultures of human erythrocytes. NET-rich supernatants were added to cultures of *P. falciparum*-infected erythrocytes and the proportion of erythrocytes presenting intracellular ring structures (A) or the proportion of infected erythrocytes (B) were determined after 48 hours. Supernatants from unstimulated neutrophils were used as control. NET-rich supernatants were treated with DNAse (C), MPO inhibitor (MPOi, 10 µg/mL) (D) or NE inhibitor (NEi, 10 µg/mL) (E) 30 minutes before adding to erythrocyte cultures and the proportion of erythrocytes presenting ring structures was determined after 48 hours as in A. *** P< 0.001 relative to controls.

We then moved to a murine model of malaria, using *P. berghei* ANKA and bone marrow-derived neutrophils from C57/BL6 mice. *P. berghei* ANKA is known to induce a severe form of cerebral malaria in susceptible C57/BL6 mice and is generally used as a model for the human form of *P. falciparum*-induced cerebral malaria. Similarly to what we found in human neutrophils, incubation of mouse neutrophils with *P. berghei*-infected red blood cells induced a significant increase in extracellular DNA that was not observed in neutrophils incubated with uninfected erythrocytes (Fig 6A). NET release by murine neutrophils was also ROS-independent since it was unaffected by either DPI (Fig 6B) or NAC treatment (S5A Fig), despite the strong ROS production induced by infected red blood cells (S5B Fig). Morphologically, NETs from murine neutrophils also stained positively for MPO and citrullinated histones, but were slightly distinct from NETs released by human neutrophils in that it showed a fiber-like structure (Fig 6C). Together, these results show that, similarly to what we described for human neutrophils in response to *P. falciparum*-infected erythrocytes, murine neutrophils release NET in response to *P. berghei* ANKA-infected erythrocytes, in a process that is independent of ROS.

**Fig 6.**
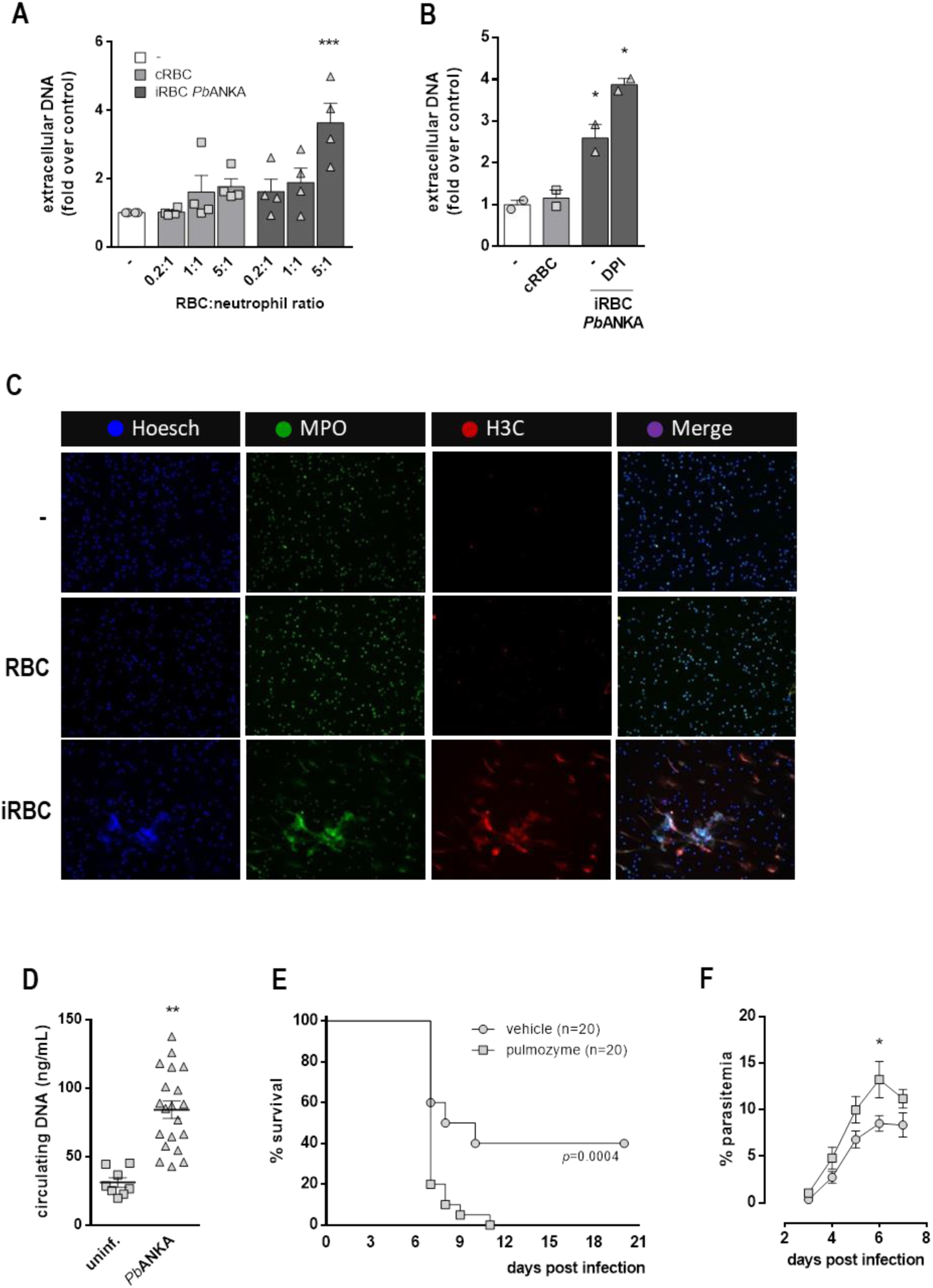
*P. berguei* ANKA-infected erythrocytes induce NETs. (A) Fluorimetric determination of NET production by murine neutrophils in the presence of *P. berguei* ANKA-infected mouse red blood cells (iRBC *Pb*ANKA) or uninfected RBC (cRBC) at varying red blood cell:neutrophil ratios. (B) Mouse neutrophils were pre-treated for 30 minutes with DPI and then incubated with *Pb*ANKA-infected RBC. NET production was determined by fluorimetry as before. Data in A and B are presented as means ± S.E.M. of the fold induction of extracellular DNA signal relative to resting neutrophils. (C) Visualization by fluorescence microscopy of NETs produced by murine neutrophils in the presence of *Pb*ANKA-infected (iRBC) or uninfected (cRBC) red blood cells. DNA is stained in blue (Hoesch), myeloperoxidase is stained in green (MPO) and citrullinated histone H3 (H3C) is stained in red. Unstimulated human neutrophils were used as controls. (D) Determination of circulating levels of DNA in plasma of *P. berguei* ANKA-infected C57BL6 mice 6 days after infection. (E) Mice were treated with DNAse (Pulmozyme, 5 mg/kg i.p., 1 hour before and every 8 hours for 6 days) and survival after *Pb*ANKA infection was monitored for 21 days. (F) Parasitemia of mice treated with DNAse (as in E) or vehicle and infected with *Pb*ANKA was monitored daily for 7 days. Data are presented as means ± S.E.M. of the percentage of infected red blood cells. * P< 0.05 and ** P<0.01 relative to untreated controls.

Finally, infection of C57/BL6 mice with *P. berghei* ANKA resulted in increased plasmatic levels of circulating DNA (Fig 6D), which corroborates with our data from S.M. patients (Fig 1B). *P. berghei* ANKA infection also resulted in sharp mortality starting at day 7 and that reached a 40% survival rate by day 10 (Fig 6E). Treatment of mice with DNAse (Pulmozyme) resulted in accelerated death, with 20% survival at day 7 and 100% mortality by day 10 post-infection (Fig 6E). Interestingly, this DNAse effect was paralleled by a significant increase in parasitemia (Fig 6F), suggesting that NET functions to restrain parasite dissemination *in vivo* in a similar fashion to what we observed *in vitro*.

## Discussion

Herein we show that *Plasmodium*-infected red blood cells release MIF that induce NET formation by human and mouse neutrophils *in vitro*. Addition of purified NET to infected erythrocyte cultures reduced the proportion of parasite-positive cells in a mechanism dependent on MPO and NE. Since MPO and NE are cytotoxic, it is possible that in malaria infection NET serves not only as a trap, but also to kill free parasites. Patients suffering from severe malaria have increased amounts of circulating DNA, paralleled by increased NE levels in plasma. To gain insight into the contribution of NET to malaria pathophysiology, we used a well described mouse model of severe malaria caused by *P. berghei* ANKA. Infected mice had higher amount of circulating DNA and treatment with DNAse increased parasitemia and accelerated mortality, supporting a role for NET in the resistance against malaria infection.

*In vitro*, NET release in response to *Plasmodium*-infected erythrocytes occurred early (starting within the first 15 minutes of stimulation), was dependent of histone citrullination by PAD4 and independent of ROS, MPO or NE. This resembles the processes described as non-lytic NET release, documented in response to *Candida albicans*, *Staphylococcus aureus* and *Escherichia coli* [32, 40, 41]. On the other hand, we found that NET release was accompanied by significant increase in extracellular LDH activity, suggestive of cell death. This LDH activity could come from neutrophils producing NET, from lysis of erythrocytes or both.

We observed that MIF, acting through CXCR4, were required to NET release induced by infected red blood cells. Recent evidences suggest that red blood cells are a major source of MIF in the bloodstream [39]. The mechanism by which MIF is released from infected erythrocytes is not well characterized. MIF might be released after red cell lysis during the parasite cycle. However, unless erythrocytes lyse immediately upon co-culture with neutrophils, only the continuous release of MIF would explain NET being triggered as soon as 15 minutes. One possible alternative is that MIF could be released within red blood cell-derived microvesicles that are continuously shed by infected erythrocytes independently of parasite cycling [42]. Another possibility is that in response to infection, red blood cells are stimulated to continuously release MIF independent of microvesicles. A previous study reported that MIF potentiates *Pseudomonas aeruginosa*-induced NET release in both humans and murine neutrophils [43]. The mechanisms and signaling pathways triggered by the MIF/CXCR4 axis that contribute to NET release require further investigations.

The protective role of NET described here contrasts with reports demonstrating the contribution of NETs to tissue damage upon experimental *Plasmodium* infection. DNAse treatment of mice, or neutrophil depletion, alleviated lung injury and resulted in increased survival of mice in a model of malaria-associated acute lung injury [27]. Moreover, neutrophil depletion has been shown to be beneficial in different studies [12, 22, 23, 44]. Most of these studies, however, use different mouse strains and *Plasmodium* species, which may account for differences in outcome. Ioannidis and cols. used the same experimental model of *P. berghei* ANKA infection of cerebral malaria susceptible C57B6 mice [12]. In their study neutrophils played a detrimental role as a significant source of CXCL10, since neutrophil depletion or CXCL10 ablation prevented C.M. development. These results can be reconciled when considering a double-edged role for neutrophils in malaria pathogenesis: NET release could be beneficial, by limiting parasite dissemination, but overt neutrophil activation would result in tissue injury that overcomes any benefit. This can explain why depleting neutrophils results in increased survival while targeting NET alone (with DNAse treatment) results in increased susceptibility in the mouse model of C.M. caused by *P. berghei* ANKA. Similar to our finding in malaria patients, a recent report showed evidences of NET formation in human patients suffering from complicated malaria which positively correlated with clinical manifestations [26]. It is possible that overt neutrophil activation is occurring in these patients, resulting in increased NET release but also increased NET-independent tissue injury, i.e. by increased proteolytic activity, cytokine release and/or increased oxidative stress. Therefore, attempts to target NET or neutrophils in malaria should be taken with caution and consider the complex interplay between both beneficial and detrimental roles played by neutrophils in malaria.

## Materials and methods

### Human studies

Prior to enrollment, written informed consent was obtained from the parents/guardians on behalf of their children after receiving a study explanation form and oral explanation from a study clinician in their native language. The protocol and study procedures were approved by the institutional review board of the National Institute of Allergy and Infectious Diseases at the US National Institutes of Health (ClinicalTrials.gov ID NCT01168271), and the Ethics Committee of the Faculty of Medicine, Pharmacy and Dentistry at the University of Bamako, Mali.

### Description of population and study site

Children aged 0-10 years of age were enrolled in the health district of Ouélessébougou. Ouélessébougou is located about 80 km south from Bamako, the capital city of Mali, and contains the district health center and a Clinical Research Center located in the community health center where studies of malaria and other infectious disease have been ongoing since 2008. The district covers 14 health sub-districts. In 2008, in the town of Ouélessébougou, the incidence rate of clinical malaria in under-5 year-olds was 1.99 episodes/child/year and the incidence rate of severe malaria as defined by WHO criteria was about 1-2% in this age group during the transmission season. Malaria is the most frequent cause of admission in the pediatric service, representing 44.9% of admissions, followed by acute respiratory infections (26.4%) and diarrhea (11.2%) [45]. Malaria transmission is highly seasonal in the study area.

### Blood Collection

Samples were collected from 23 children with severe malaria (from the febrile hospitalization cohort) and 42 participants with mild malaria (from the longitudinal under-5 cohort, matched for age). Of the 23 severe malaria cases, 8 had cerebral malaria while the remaining had severe anemia or prostration. Venous blood was drawn in EDTA tubes, and plasma was prepared by centrifuging for 10 min at 1500g. Plasma was aliquoted and stored at −80°C.

### ELISA

ELISA kits used were Neutrophil Elastase (Abcam 119553, plasma dilution 1:500; standard range, 0.16-10 ng/ml) and Cell Death ELISA (Roche, 11774425001, plasma dilution 1:2) which estimates cytoplasmic histone-associated DNA fragments (mono-and oligonucleosomes, no standard range). For the ordinal variables, differences between groups were calculated using the non-parametric Mann-Whitney test.

### Neutrophil purification

Human neutrophils were isolated from peripheral blood using a histopaque 1077 density gradient as previously described [46]. Erythrocytes were lysed with ACK solution and the pellet containing neutrophils was washed in HBSS and ressuspended in cold RPMI 1640 medium. Isolated neutrophils were routinely ≥ 95% pure and >99% viable. Murine neutrophils were isolated from the bone marrow of C57/BL6 mice by percoll density gradient as described [47]. Isolated neutrophils were resuspended in cold RPMI 1640 medium. Purity was routinely ≥ 95% pure and viability >99%.

### Neutrophil treatment

To evaluate the participation of ROS in NET production, neutrophils were pre-treated with diphenyleneiodonium chloride (DPI, Sigma-Aldrich, 10 µg/mL) or N-Acetyl-L-cysteine (NAC, Sigma-Aldrich, 10 µM). Neutrophils were pretreated with pharmacological inhibitors to MPO (MPOi, Santa Cruz Biotechnology, 10 µg/mL), neutrophil elastase (NEi, Santa Cruz Biotechnology, 10 µg/mL), PDA4 (Cl-amidine, Cayman Chemical, 12 µM), CXCR4 (AMD3100, Sigma-Aldrich, 100 ng/ml), MIF (ISO-1, Abcam, 50μM), JNK (SP600125, Sigma-Aldrich, 40 μM), p38 MAPK (SB239063, Sigma-Aldrich, 20 μM), and phagocytosis (cytochalasin-D, Sigma-Aldrich, 8 μM). Finally, neutrophils were also treated with blocking antibodies to MIF (kindly provided by Dr. R. Bucala, 20μg/ml), CD18 (20μg/ml, Abcam), CD36 (20μg/ml, Abcam) or ICAM-1(20μg/ml; R&D Systems), or the appropriate isotype control IgG (20 μg/ml, Abcam). All inhibitors and antibodies were added to neutrophil cultures 30 minutes before stimulation.

### Parasite cultures

*Plasmodium falciparum* W2 strain was cultured in human A+ type erythrocytes at 37°C under controlled gas atmosphere (5% CO_2_, 5% O_2_, 90% N_2_), in RPMI supplemented with 20 mM HEPES, 22 mM glucose, 0.3 mM hypoxanthine, 0.5 % albumax II and 20 µg/mL of gentamycin [48]. Culture parasitemia was determined daily through thick blood smear stained with Diff-Quick and maintained around 2% at a 4 to 5% hematocrit. Parasitemia (number of infected RBC per 100 RBCs) was determined by counting at least 500 cells. In a selected experiment, supernatant from infected cultures was collected and immediately added to neutrophil cultures. Supernatant from uninfected erythrocytes was used as control.

### Mature trophozoites purification

Mature trophozoites were isolated by percoll/sorbitol gradient as described previously [49]. Briefly, cultures of infected erythrocytes with at least 5% parasitemia were centrifuged at 900g for 15 min at room temperature. Pellet was resuspended in fresh RPMI to reach a 20% hematocrit and gently poured on top of a 40%, 70% and 90% Percoll/sorbitol gradient. After centrifugation the brown band formed between the 40% and 70% layers was harvested and suspensions of synchronized trophozoites (>90% of purity) were used to stimulate neutrophils.

### Fluorimetric quantification of NETs

Neutrophils (2×10^5^ cells) were stimulated with *P. falciparum*-infected erythrocytes at varying neutrophil:erythrocyte ratios. After incubation, ECOR1 and HIINDIII restriction enzymes (20 units/mL each) were added and incubated for 30 min at 37°C. Samples were then centrifuged and supernatants collected. DNA concentration in the supernatants (referred to as NETs) was determined using Picogreen dsDNA kit (Invitrogen) according to the manufacturer’s instructions. Uninfected erythrocytes from the same blood type were used as control.

### Visualization of NETs by immunofluorescence

Neutrophils (2×10^5^) were allowed to adhere onto 0.001% poly-L-lysine (Sigma) coated glass coverslips. Neutrophil were then stimulated with 1×10^6^ *P. falciparum*-infected erythrocytes for 3 h. Cells were fixed with 4% paraformaldehyde for 15 min at room temperature. After extensive wash in PBS, unspecific binding sites were blocked with 3% BSA and cells were incubated with primary anti-myeloperoxidase (1:1000, Abcam), anti-elastase (1:1000, Abcam), or anti-citrullinated histone H3 (1:1000, Abcam) antibodies, followed by the appropriate secondary fluorescent antibodies (1:4000). DNA was counterstained with Hoesch. Images were acquired using a Leica confocal microscope under 40X and 100X magnification.

### Quantification of ROS production

ROS production was measured using a fluorimetric assay based on the oxidation of the CM-H2DCFDA probe (Molecular Probes) following the manufacturer’s instructions. Briefly, 2×10^5^ neutrophils and 10^6^ infected erythrocytes were mixed with 2 μM of CM-H2DCFDA probe in a 96 well plate. Fluorescence was monitored every 10 min for 30 min. Uninfected erythrocytes were used as controls. The same culture and stimulation procedure was carried out for the visualization of ROS production under the microscope. Images were acquired using a Leica DMI6000 fluorescence microscope under 20x magnification after 1 hour of stimulation.

### Parasite invasion and growth assays

NET-rich supernatant was obtained from human neutrophils cultured with *P. falciparum*-infected erythrocytes at a 1:10 ratio for 3 hours. Cultures were centrifuged and NET-rich supernatant was collected for immediate use. Supernatant obtained from neutrophils incubated with uninfected red blood cells was used as control. Infected erythrocytes at 2% parasitemia were seeded in a 96-well plate in RPMI supplemented with 10% FCS to reach a 5% hematocrit. NET-rich supernatants were added to the erythrocyte cultures which were incubated at 37°C for 24 h. Parasite invasion was estimated by counting the number of new intracellular ring forms in a thick blood smear stained with Diff-Quick. Invasion was expressed as the percentage of erythrocytes showing ring forms of the parasite. Additionally, cultures were allowed to proceed for up to 48 h to analyze intracellular parasite growth. The number of infected erythrocytes, including all parasite forms, was determined and expressed as the percentage of infected red blood cells (iRBC).

### Westernblot

Whole-cell lysates were extracted by RIPA buffer and cleared by centrifugation at 15000×g for 15 min at 4°C prior to boiling in Laemmli buffer. Western blots were performed using standard molecular biology techniques and membranes were developed using Super Signal West Femto Maximum Sensitivity Substrate (Thermo Scientific). Blot images were acquired in a ChemiDoc XRS system (BioRad). Antibodies used were anti-p-JNK (Cell Signaling Technologies), anti-p-p38 (BD Biosciences), anti-p AKT (Cell Signaling Technologies), and anti-β-actin (Millipore). All primary antibodies were diluted 1:1000 in TBS-T.

### In vivo assays

All animal procedures were approved by the Institution Ethics Committee (CEUA protocol number XXX). C57BL6 mice were treated intravenously with either vehicle (0.9% NaCl sterile saline) or Pulmozyme (5 mg/kg, Roche) 1 hour before infection. Pulmozyme treatment was continued every 8 hours for 6 days. Mice were infected with 1×10^5^ *P. berghei* ANKA. Mice were monitored daily for clinical signs of cerebral malaria and blood samples were collected daily for parasitemia determination.

### Statistical analysis

Data are presented as means ± S.E.M. of at least 3 independent experiments. All statistical analyses were performed using GraphPad Prism 6.0 for windows. One-way ANOVA was used for comparisons among multiple groups. Survival analysis was carried out using the built-in Prism survival analysis. Paired Student t-test was used to compare differences between cultures in the presence and absence of NET-rich supernatant. Differences with P˂ 0.05 were considered as statistically significant.

## Acknowledgements

The authors thank Prof. Richard Bucala and Dr. Lin Leng (Yale School of Medicine) for providing anti-MIF neutralizing monoclonal antibody ascite, and members of the Bozza Lab for helpful discussions. This work was financially supported by Conselho Nacional de Pesquisa (CNPq), Coordenação de Aperfeiçoamento de Pessoal de Nível Superior (CAPES) and Fundação de Amparo à Pesquisa do Rio de Janeiro (FAPERJ).

## Supporting information

**S1 Fig.**
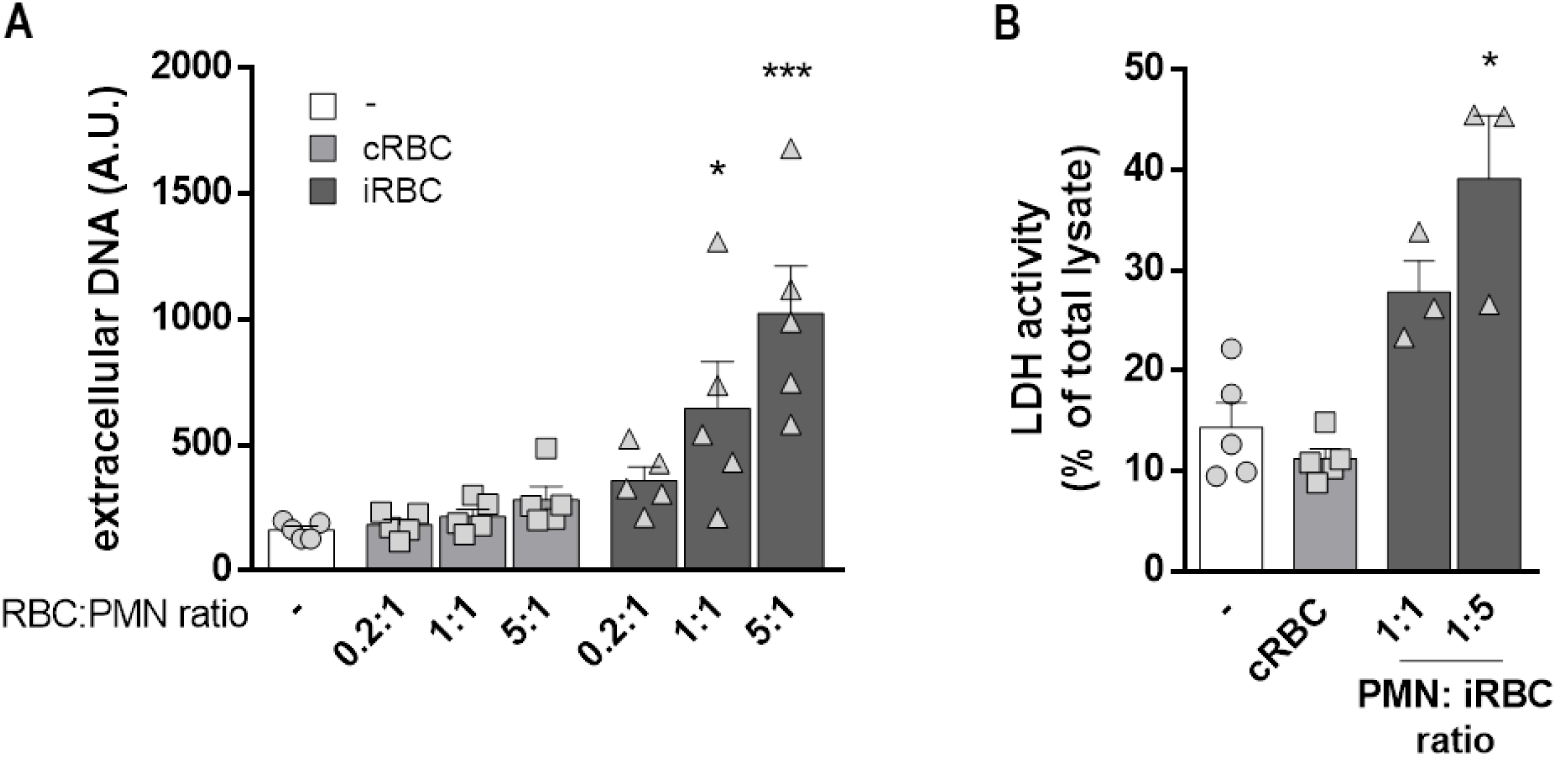
(A) Fluorimetric determination of NET production by human neutrophils in the presence of *P. falciparum*-infected red blood cells (iRBC) or uninfected RBC (cRBC) at varying red blood cell:neutrophil ratios and represented as means ± S.E.M. of the extracellular DNA fluorescence signal (in arbitrary units). (B) Determination of lactate dehydrogenase (LDH) activity in culture supernatants of human neutrophils incubated with infected red blood cells (iRBC) at two different neutrophil:red blood cell ratios for 3 hours. Uninfected red blood cells (cRBC) were used as controls. LDH activity in culture supernatants was compared to the total intracellular LDH activity as determined in neutrophil cell lysates. * P< 0.05 and *** P< 0.001 relative to unstimulated neutrophils.

**S2 Fig.**
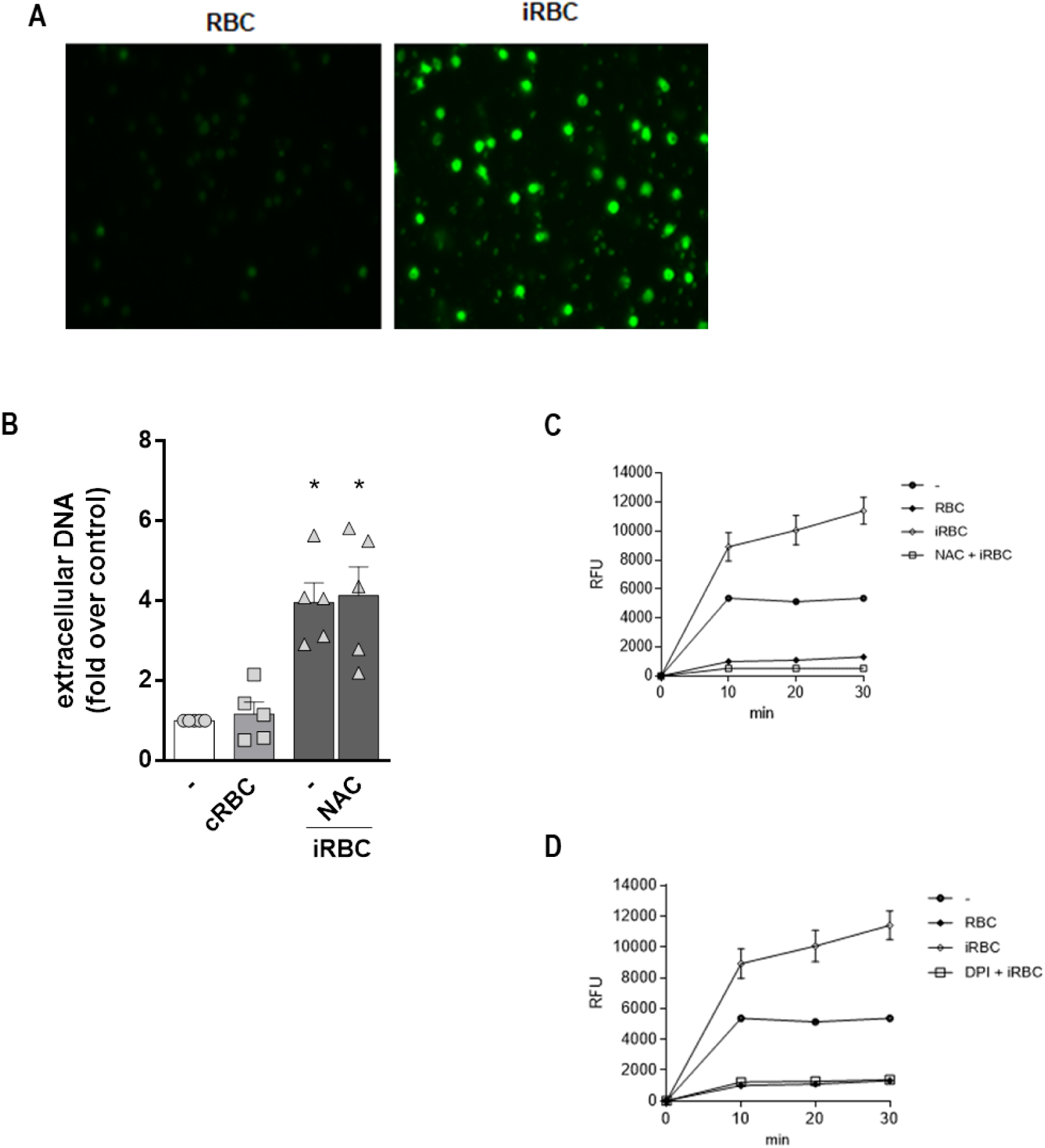
(A) Representative fluorescence images of ROS production by human neutrophils incubated with infected (iRBC) or uninfected red blood cells (RBC) at a 1:5 ratio in the presence of the ROS-sensitive CM-H2DCFDA probe. (B) Human neutrophils were treated with NAC (10 µM) for 30 minutes and then incubated with *P. falciparum*-infected red blood cells (iRBC). NET production was determined by fluorimetry. Uninfected red blood cells (cRBC) were used as control. Data are presented as means ± S.E.M. of the fold induction of extracellular DNA signal relative to resting neutrophils. (C and D) Kinetics of ROS production by human neutrophils incubated with infected red blood cells (iRBC) and treated or not with antioxidants DPI (C) or NAC (D). ROS production was evaluated by fluorimetry every 10 minutes for 30 minutes in the presence of CM-H2DCFDA.

**S3 Fig.**
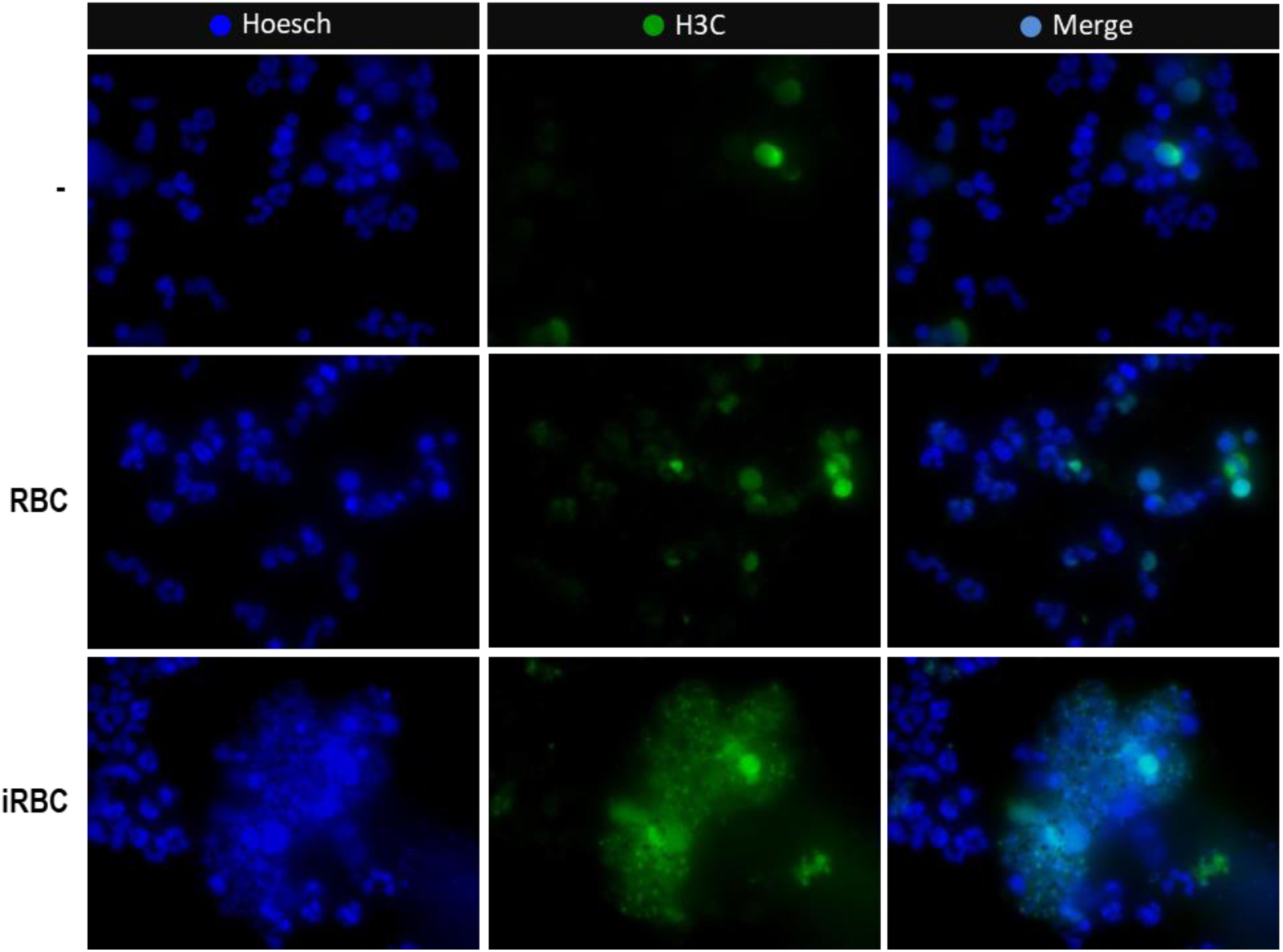
Representative immunofluorescence images of human neutrophils incubated with *P. falciparum*-infected (iRBC) or uninfected (cRBC) red blood cells at a 1:5 ratio for 3 hours and stained for DNA (blue) and citrullinated histone H3 (green). Unstimulated neutrophils were used as controls.

**S4 Fig.**
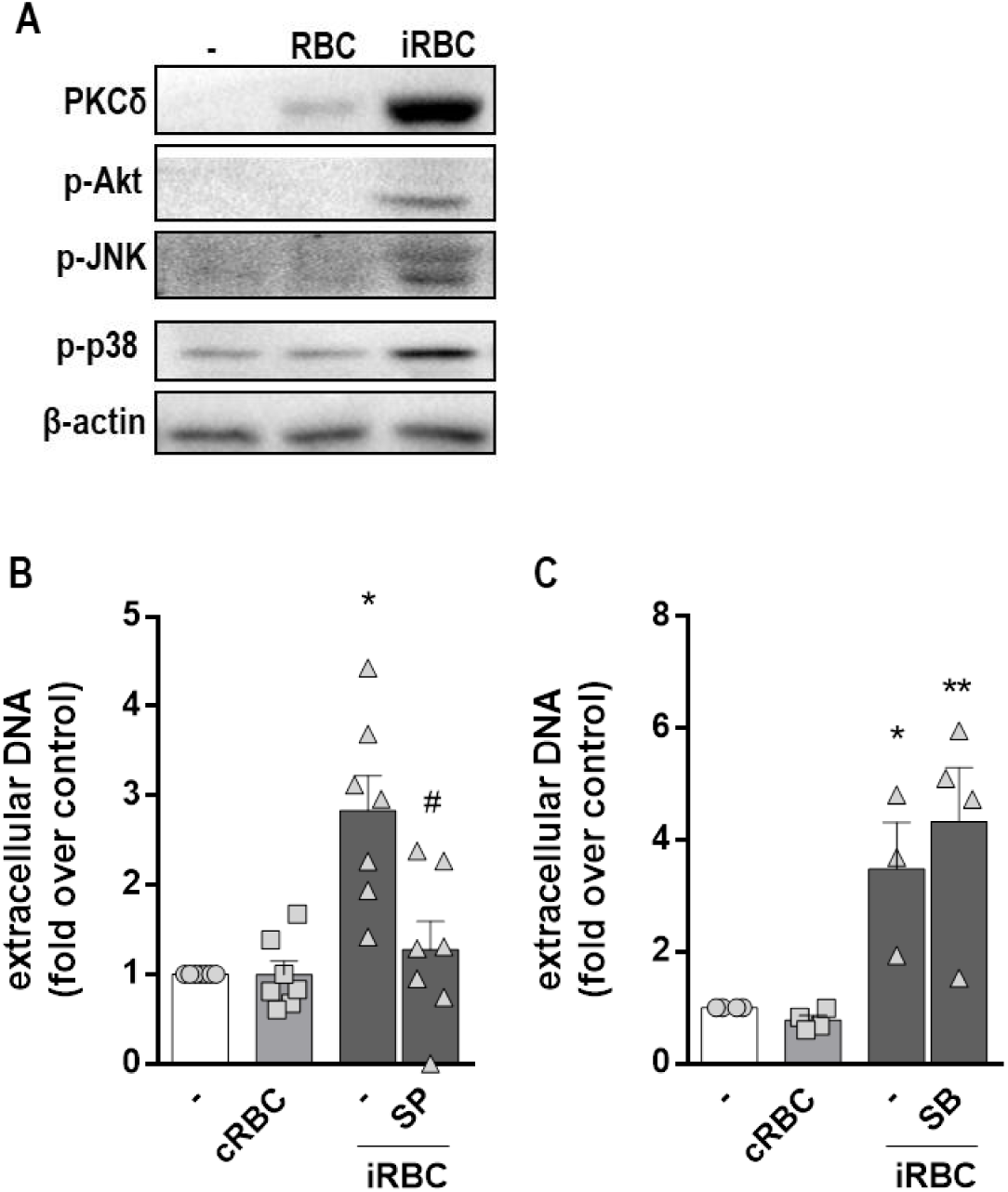
(A) Representative westernblot images of total cell extracts of neutrophils incubated with infected (iRBC) or uninfected (cRBC) red blood cells at a 1:5 ratio. Westernblot was used for the detection of total PKCδ and phosphorylated Akt (p-Akt), JNK (p-JNK) and p38 (p-p38). β-actin was used as loading control. Unstimulated neutrophils were used as controls. (B and C) Human neutrophils were treated with SB239063 (SB, 20 μM) (B) or SP600125 (SP, 40 μM) (C) for 30 minutes and then incubated with *P. falciparum*-infected red blood cells (iRBC) at a 1:5 ratio for 3 hours. NET production was determined by fluorimetry. Uninfected red blood cells (cRBC) were used as control. Data are presented as means ± S.E.M. of the fold induction of extracellular DNA signal relative to resting neutrophils. * P< 0.05 and ** P< 0.01 relative to controls incubated with cRBC, # P< 0.01 relative to untreated control.

**S5 Fig.**
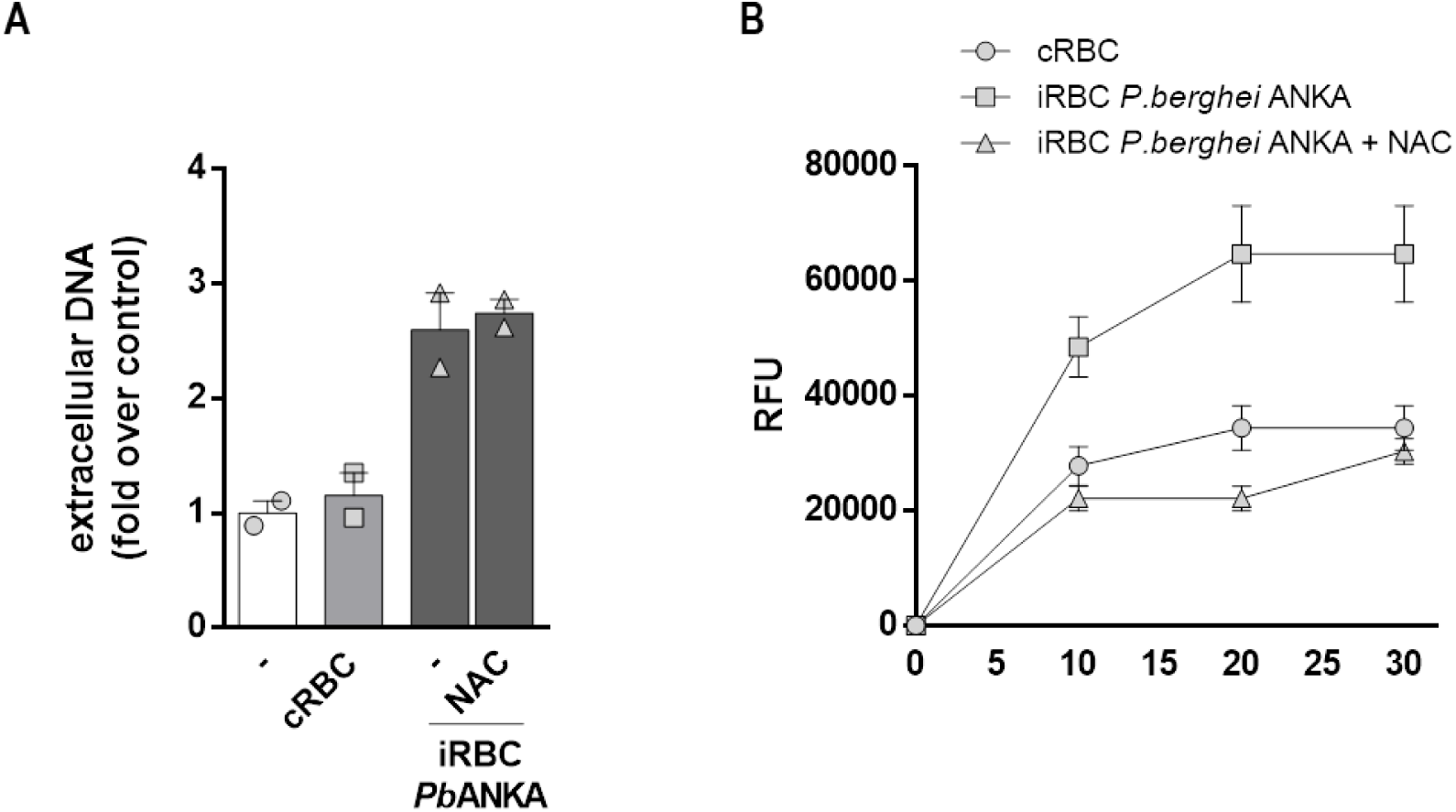
(A) Murine neutrophils were treated with NAC (10 µM) for 30 minutes and then incubated with *P. berguei* ANKA-infected red blood cells (iRBC). NET production was determined by fluorimetry. Uninfected red blood cells (cRBC) were used as control. Data are presented as means ± S.E.M. of the fold induction of extracellular DNA signal relative to resting neutrophils. (B) Kinetics of ROS production by murine neutrophils incubated with infected red blood cells (iRBC) and treated or not with NAC. ROS production was evaluated by fluorimetry every 10 minutes for 30 minutes in the presence of CM-H2DCFDA.

